# Phosphatase PP2C6 negatively regulates phosphorylation status of Pti1b kinase, a regulator of flagellin-triggered immunity in tomato

**DOI:** 10.1101/613612

**Authors:** Fabian Giska, Gregory B. Martin

## Abstract

Plant immune responses, including the production of reactive oxygen species (ROS), are triggered when pattern recognition receptors (PRR) become activated upon detection of microbe-associated molecular patterns (MAMPs). Receptor-like cytoplasmic kinases are key components of PRR-dependent signaling pathways. In tomato two such kinases, Pti1a and Pti1b, are important positive regulators of the plant immune response. However, it is unknown how these kinases control plant immunity at the molecular level, and how their activity is regulated. To investigate these issues, we used mass spectrometry to search for interactors of Pti1b in *Nicotiana benthamiana* leaves and identified a protein phosphatase, PP2C6. An *in vitro* pull-down assay and *in vivo* split luciferase complementation assay verified this interaction. Pti1b was found to autophosphorylate on threonine-233 and this phosphorylation was abolished in the presence of PP2C6. An arginine-to-cysteine substitution at position 240 in the Arabidopsis MARIS kinase was previously reported to convert it into a constitutive-active form. The analogous substitution in Pti1b made it resistant to PP2C6 phosphatase activity, although it still interacted with PP2C6. Treatment of *N. benthamiana* leaves with the MAMP flg22 induced threonine phosphorylation of Pti1b. Expression of PP2C6, but not a phosphatase-inactive variant of this protein, in *N. benthamiana* leaves greatly reduced ROS production in response to treatment with MAMPs flg22 or csp22. The results indicate that PP2C6 acts as a negative regulator by dephosphorylating the Pti1b kinase, thereby interfering with its ability to activate plant immune responses.

## Introduction

Plants exist in an environment containing large numbers of diverse microorganisms, some of them detrimental to plant health. Consequently, their survival depends on their ability to specifically recognize and respond to pathogenic microbes. This recognition is enabled by the function of plasma membrane-localized pattern recognition receptors (PRRs), which recognize microbe-associated molecular patterns (MAMPs) (1-3). The best understood PRR is FLS2, which perceives the peptide flg22, a part of the flagellin protein, which forms the bacterial flagellum (4,5). Two PRRs that are present only in solanaceous species, including tomato, are FLS3, which recognizes a flagellin-derived peptide, flgII-28, and CORE, which detects a MAMP in the bacterial cold shock protein, csp22 (6,7). The binding of the extracellular part of the receptor to the appropriate ligand activates the cytoplasmic kinase domain of the receptor (8).

Activated PRRs induce a defense response referred to as pattern triggered immunity (PTI). PTI is associated with the generation of reactive oxygen species (ROS), activation of mitogen-activated protein kinases (MAPKs), transcriptional reprogramming, production of antimicrobial molecules, and the reinforcement of the cell wall, among other processes (9-12). Receptor-like cytoplasmic kinases (RLCK) play a key role in linking PRRs to downstream signaling components (13). RLCKs constitute a large family of proteins consisting primarily of a serine-threonine kinase domain but some members also possess EGF, LRR, WD40 and transmembrane domains (13).

The *Arabidopsis thaliana* genome encodes 179 RLCKs divided into 17 subfamilies: RLCK-II and from RLCK-IV to RLCK-XIX. The majority of RLCKs which are known to have a role in plant immunity belong to sub-family VII. For example, AtBIK1, a member of sub-family VII, is activated upon phosphorylation by FLS2 after its perception of flg22 (13,14). The activated BIK1 induces generation of ROS by phosphorylating the NADPH oxidase, RBOHD (11,15). Similarly, the activated PRR chitin elicitor receptor kinase 1 (CERK1) phosphorylates the RLCK protein PBL27, which then directly activates an immunity-associated MAPK kinase cascade (16). Some RLCKs play a negative regulatory role such as PBL13 whose activation reduces generation of ROS in Arabidopsis and hinders resistance against the bacterial pathogen *Pseudomonas syringae* pv. tomato (17).

RLCKs also play important roles in regulating plant growth, development and reproduction. One such RLCK is MARIS, which belongs to sub-family VIII. MARIS was identified in a forward genetic screen as a suppressor of the *anx1/anx2* mutations (18). This double mutation disrupts cell wall integrity (CWI)-signaling and results in a burst of the pollen tube after pollen germination. The *MARIS* allele identified in the screen contains a mutation that introduces an arginine-to-cysteine substitution at position 240 (R240C) in the activation loop of the kinase which rescues the pollen tube bursting phenotype of the *anx1/anx2* mutant (18). This and other observations suggested that MARIS acts downstream of the ANX1/ANX2 receptors in CWI signaling and that the R240C variant is a constitutively-active form of MARIS (18), although the molecular basis for this constitutive activity is unknown.

Defense responses in plants need to be tightly regulated to prevent damage to plant tissue. Since immune responses are often activated by protein kinases, protein phosphatases are natural candidates as a negative regulators of plant responses to pathogen attack. Indeed a few phosphatases are known to function as regulators of PRRs, immunity-associated RLCKs, and MAPKs. The kinase associated protein phosphatase (KAPP), for example, interacts with FLS2 and KAPP overexpression blocks flg22-dependent signaling (19). The rice PRR XA21 which confers resistance to the bacterial pathogen *Xanthomonas oryzae* pv. oryzae, interacts with XB15 which belongs to the protein phosphatase 2C (PP2C) family (20). XB15 dephosphorylates the autophosphorylated kinase domain of XA21. Rice plants with a mutation in the *Xb15* gene show symptoms of activated defense responses whereas plants overexpressing *Xb15 are* more susceptible to *X. o.* pv. oryzae (20). In *Nicotiana benthamiana*, virus-induced gene silencing of a gene encoding a catalytic subunit of protein phosphatase 2A results in activation of plant defense responses but the mechanism is not known (21). In Arabidopsis, the PRR co-receptor BAK1 interacts with a phosphatase PP2A holoenzyme. PP2A also negatively controls phosphorylation status of BAK1 thereby interfering with subsequent PTI signaling (22). The RLCK BIK1 is known to interact with phosphatase PP2C38. The phosphatase dephosphorylates BIK1 and impedes its ability to phosphorylate the NADPH oxidase RBOHB causing impaired ROS production and decreased stomatal-mediated immune responses (23). Finally, some MAPK phosphatases are known to negatively regulate aspects of PTI including ROS production (24,25).

In tomato, two highly similar kinases referred as Pti1a (Pto interactor 1a) and Pti1b play a role in pattern-triggered immunity against *P. s.* pv. tomato (26). Like MARIS, Pti1a and Pti1b are members of RLCK family VIII and consist primarily of a serine-threonine protein kinase domain (26). We previously generated and characterized tomato plants carrying a hairpin (hp)-*Pti1* construct which consequently have reduced expression of both *Pti1a* and *Pti1*b. The hpPti1 transgenic plants are impaired in ROS production in response to treatment with MAMPs flg22 or flgII-28, whereas activation of MAPK kinases is unaffected (26). Importantly, the hpPti1 plants are significantly more susceptible to *P. s*. pv. tomato. Pti1a was previously shown to intramolecularly autophosphorylate on threonine-233 in the activation loop of the kinase; however, the possible role of this phosphorylation in Pti1a-mediated PTI signaling is unknown (27,28).

Homologs of the Pti1 kinases exist in many other plant species. In Arabidopsis it was shown that Pti1-2 and Pti1-4 play a role in oxidative stress (29,30), and the kinase activity of Pti1-2 is induced by flg22 (29). Arabidopsis Pti1-1 and Pti1-2 proteins, but not Pti1-3 or Pti1-4, show strong autophosphorylation activity (29). In cucumber, a Pti1 homolog CsPti1-L is a positive regulator of plant immunity and salt tolerance, and also exhibits autophosphorylation activity (31). In contrast, it was shown in rice that OsPti1a acts as a negative regulator of basal defense against *X. o*. pv. oryzae and *Magnaporthe oryzae* (32,33). As with the tomato Pti1a kinase, threonine-233 of OsPti1a undergoes autophosphorylation *in vitro* and additionally is phosphorylated by the OsOxi1 kinase (33). The authors postulate that the negative regulatory function of OsPti1a might be abolished by OsOxi1-dependent phosphorylation of OsPti1a on threonine 233 (33). An OsPti1a(T233A) variant is unable to restore basal resistance against *X. o.* pv. oryzae in an *ospti1a* mutant background in comparison to OsPti1 wildtype (33).

Although much is known about the enzymatic activity and upstream activators of Pti1-like kinases, little is known about their downstream substrates or associated proteins that might regulate their autophosphorylation status. To find potential substrates and regulators of the Pti1b kinase we used a combined FLAG co-immunoprecipitation and Strep pull-down approach coupled with mass spectrometry. We focused our research on Pti1b because previous RNA-Seq analyses indicated that expression of the *Pti1b* gene and not the *Pti1a* gene is induced by MAMPs csp22 and flgII-28 and infection by *P. syringae* pv. tomato, suggesting it might play a more prominent role in PTI (34). We found that Pti1b expressed in *N. benthamiana* leaves co-purifies with a phosphatase of the PP2C family (NbPP2C6). We verified this interaction using additional assays and investigated the role of the phosphatase in regulating Pti1b. Our observations indicate that PP2C6 dephosphorylates Pti1b and acts as a negative regulator of PTI signaling.

## Experimental procedures

### DNA cloning

The *AtPIP2a* gene was transferred from plasmid pCambia-AtPIP2a-cLuc (kind gift from G. Coaker) into the pJLSmart Gateway entry vector (40) and then moved into the pGWB-SF destination plasmid that enables protein expression in fusion with a 2xStrep-FLAG-at the C terminus of the protein. pGWB-SF plasmid was obtained by cloning a 2xStrep sequence into pGWB411 vector (41). *Pti1b*(K96N)-pJLSmart and *Pti1b*(R234C)-pJLSmart were generated with PCR-based site-directed mutagenesis with construct Pti1b-pJLSmart (26) as the template. *Pti1b, Pti1b*(K96N) and *Pti1b*(R234C) genes were then transferred from pJLSmart into the pGWB-SF destination plasmid. For protein expression in *E. coli*, the *Pti1b* gene was amplified from Pti1b-pJLSmart with primers that enabled cloning into the pET30a+ vector using restriction enzyme *Nde*I and *Not*I. *Pti1b(*K96N), *Pti1b*(S232A), *Pti1b*(T233A), *Pti1b*(S232A,T233A), *Pti1b*(T233D), *Pti1b*(R234C) and *Pti1b*(T233A,R234C) variants were generated with PCR-based site-directed mutagenesis. The appropriate Pti1b variant in the pET30a+ plasmid was used as the template for the PCR. The *PP2C6* gene was amplified from tomato cDNA and cloned into pASK3 plasmid using restriction enzymes *Xba*I and *Afe*I. The *PP2C6*(NN) variant was generated with PCR-based site-directed mutagenesis with *PP2C6*-pASK3 as the template. For ROS assays in *N. benthamiana* PP2C6 variants were amplified from pASK3 plasmids and cloned into pJLTRBO (42) with restriction enzymes *Not*I and *Avr*II. Similarly, the *GFP* gene was amplified from plasmid pGWB505 and cloned into the pJLTRBO plasmid using the same restriction enzymes. To carry out the split luciferase complementation assay (SLCA) *Pti1b* and *PP2C6 genes* were amplified with PCR and cloned into pCambia-nLuc and pCambia-cLuc respectively using restriction enzymes *Kpn*I and *Sal*I. All primers used in this study are listed in Supplementary Tables S2 and S3.

### Bacteria growth and infiltration

*Agrobacterium tumefaciens* GV3101 was grown on LB medium with appropriate antibiotics for 48 hours at 30°C. The bacterial cells were collected and suspended in buffer containing: 10 mM MES, 10 mM MgCl2, 200 *µ*M acetosyringone, pH 5.7. For agroinfiltration into *N. benthamiana* leaves, bacteria were diluted to OD 1.0, and incubated for 3 hours at room temperature. *Nicotiana benthamiana* leaves were infiltrated with a needleless syringe.

### Protein expression and purification

For mass spectrometry analysis, proteins in fusion with 2xStrep-FLAG were purified with Anti-FLAG M2 Magnetic Beads (Sigma Co.). Purification was carried out according to the manufacturer’s protocol with minor modifications. Specifically, 5 g of *N. benthamiana* leaf tissue was ground in liquid nitrogen and dissolved in 7.5 ml of extraction buffer containing: 150 nM Tris-Cl, 150 mM NaCl, 5 mM DTT, 10 mM MnCl2, 1% [v/v] Triton X-100, PhosSTOP™ - phosphatase inhibitor tablets (Sigma), cOmplete™, Mini, EDTA-free Protease Inhibitor Cocktail (Roche Co.), pH 7.4). After 30 min incubation on a rotator at 4°C the protein samples were centrifuged at 18,000x*g* for 30 min to remove plant debris and the protein extract was filtered through Miracloth filter paper (Calbiochem Co.). Next, 20 *µ*l of previously washed anti-FLAG magnetic beads in an amount equivalent to the 20 *µ*l of magnetic bead suspension was added to the protein extract. The sample was incubated on a rotator for 1 hour at 4°C and the magnetic beads were then collected using a magnetic rack. The beads were washed 3 times with 200 *µ*l extraction buffer and eluted with 100 *µ*l of elution buffer containing: 100 mM Tris-Cl, 150 mM NaCl, 5mM DTT, 10 mM MnCl2, pH 8.0) and 450 ng/*µ*l 3xFLAG peptide. After elution, MagStrep “type 3” XT beads (https://www.iba-lifesciences.com) in an amount equivalent to the 10 *µ*l of magnetic bead suspension was added to the sample. After 1.5 hours of incubation on a rotator at 4°C the beads were washed two times with 200 *µ*l of washing buffer containing: 100 mM Tris-Cl, 150 mM NaCl, 5mM DTT, pH 8.0). The proteins bound to the beads were then eluted by boiling in Laemmli buffer.

To purify 6xHis-tagged proteins a frozen pellet of *E. coli* strain DE3(pLys)Rosetta was thawed in buffer containing: 50 mM Tris-Cl, 1 M NaCl_2,_ 10mM Imidazole, 5% glycerol, cOmplete™ Mini EDTA-free Protease Inhibitor Cocktail (Roche Co.), 1mg/ml lysozyme, pH8.0). After sonication the protein extract was centrifuged at 18,000x*g* for 30 min to remove bacterial debris. The supernatant was incubated on a rotator with Ni-NTA Agarose (Qiagen Co.) for 1.5 hours at 4°C. Next, the sample was loaded onto a column and after liquid removal the resin was washed three times with 5 ml of buffer containing: 100 mM Tris-Cl, 1 M NaCl_2,_ 10% glycerol, 10 mM imidazole, pH 8.0). After washing, the protein bound to the resin was eluted with buffer containing: 50 mM Tris-Cl, 1 M NaCl_2,_ 10% glycerol, 250 mM imidazole, pH 8.0). For purification of Strep-tagged proteins, frozen *E. coli* strain DE3(pLys)Rosetta cells were thawed in buffer containing: 150 mM Tris-Cl, 150 mM NaCl_2_, 5mM DTT, cOmplete™ Mini EDTA-free Protease Inhibitor Cocktail (Roche) pH 8.0). After sonication, the protein extract was centrifuged 18,000x*g* for 30 min to remove bacterial debris. The supernatant was incubated on a rotator with Strep-Tactin Macroprep resin (IBA Co.) for 1.5 hours at 4°C. Next, the resin was washed 3 times with 5 ml of buffer: 150 mM Tris-Cl, 150 mM NaCl_2_, 5mM DTT pH 8.0, next the proteins bound to the resin were eluted with buffer containing: 150 mM Tris-Cl, 150 mM NaCl_2_, 5mM DTT, 2.5 mM biotin, pH 8.0).

### Western blotting

Protein samples were separated in 10% bisacrylamide gels at 200 V. Detection of phosphorylated proteins was performed in blocking solution (5% BSA, 0.1% [v/v] Tween-20) with anti-phosphothreonine antibodies (9381, Cell Signaling) and with anti-rabbit-HRP (W4011, Promega). Detection of tagged proteins was carried out in blocking solution (5% fat-free milk, 0.5% [v/v] Tween-20). Other antibodies used include: StrepMAB-Classic, HRP conjugate (2-1509-001, IBA), anti-HA High Affinity (11867423001, Roche), anti-polyHistidine antibody (H1029, Sigma) ANTI-FLAG® M2 (F1804, Sigma), anti-first 258 amino acids of luciferase (NB600, Novus), anti-C-terminal part of luciferase (sc-74548, Santa Cruz Biotechnology), anti-GFP (11814460001, Roche), anti-rat-HRP (sc-2006, Santa Cruz Biotechnology), and anti-mouse-HRP (sc2005, Santa Cruz Biotechnology).

### Mass spectrometry analysis

Peptide mixtures were analyzed by LC-MS-MS/MS (liquid chromatography coupled to tandem mass spectrometry) using Nano-Acquity (Waters) LC system and LTQ-FT-Orbitrap mass spectrometer (Thermo Electron Corp., San Jose, CA). Prior to the analysis, proteins were subjected to a standard “in-solution digestion” procedure during which proteins were reduced with 50 mM TCEP (for 60 min at 60°C), alkylated with 200 mM MMTS (45 min at room temperature) and digested overnight with trypsin (sequencing grade modified Trypsin - Promega V5111).

The peptide mixture was applied to RP-18 precolumn (nanoACQUITY Symmetry® C18 – Waters 186003514) using water containing 0.1% TFA as a mobile phase and then transferred to a nano-HPLC RP-18 column (nanoACQUITY BEH C18 - Waters 186003545) using an acetonitrile gradient (5% - 35% AcN) for 180 min in the presence of 0.05% formic acid with a flowrate of 250 nl/min. Column outlet was directly coupled to the electrospray ion source of the spectrometer working in the regime of data dependent MS to MS/MS switch. A blank run to ensure lack of cross contamination from previous samples preceded each analysis. The data acquired were processed by Mascot Distiller followed by Mascot Search (Matrix Science, London, UK, on-site license) against the database ‘NBenthLuteoviridaecRAPBSAProtAmod’ is accessible at: ftp://ftp.solgenomics.net/proteomics/Nicotiana_benthamiana/Cilia_lab/proteomics_db/ (Stacy DeBlasio and Michelle Cilia, 2014 personal communication). Search parameters for precursor and product ions mass tolerance were 30 ppm and 0.1 Da, respectively, enzyme specificity: trypsin, missed cleavage sites allowed: 1, fixed modification of cysteine by methylthio and variable modification of methionine oxidation. Peptides with Mascot Score exceeding the threshold value corresponding to <5% expectation value, calculated by Mascot procedure, were considered to be positively identified. Detected phosphorylated peptides were manually inspected.

### Kinase assay

For kinase assays, the proteins were incubated at room temperature in buffer containing: 150 mM HEPES, 150 mM NaCl_2_ 10 mM MgCl_2_, 10 mM MnCl_2_, 2 mM DTT, 20 *µ*M ATP and 1 *µ*Ci of [^32^P]γ-ATP (Perkin Elmer Co.) pH 8.0. The total volume of the reaction mixture was 20 *µ*l. The reaction was terminated by adding Laemmli buffer and boiling for 5 min. The proteins were separated on 10% polyacrylamide gels and visualized by autoradiography. To confirm protein loading the gels were stained with Coomassie Blue.

### Strep-tag pull-down assay

To decrease nonspecific interactions between MagStrep “type 3” XT beads and proteins the beads were incubated in 100 *µ*l of buffer containing: 150 mM HEPES, 150 mM NaCl_2_ 10 mM MgCl_2_, 10 mM MnCl_2_, 2 mM DTT, 20 *µ*M ATP, 0.1 *µ*g/*µ*l BSA, 1% Tween, pH 8.0 on a rotator for one hour at 4°C. Next 30 *µ*g of 6xHis-tagged Pti1b-variants and PPc6 in fusion with Strep-tag were added to the solution with the beads. Samples were incubated for 2 hours on a rotator at 4°C and the beads were collected using a magnetic rack and washed 2 times with 300 *µ*l of washing buffer containing: 150 mM HEPES, 150 mM NaCl_2_ 10 mM MgCl_2_, 10 mM MnCl_2_, 2 mM DTT, 20 *µ*M ATP, 1% Tween, pH 8.0. Next, the beads were resuspended again in the washing buffer and transferred to a new Eppendorf tube. The beads were collected with a magnetic rack and the proteins were eluted by boiling in Laemmli buffer. The obtained samples were separated on 10% polyacrylamide gels and transferred to a PVDF membrane. The presence of 6xHis-tagged proteins and Strep-tagged proteins was tested with 6xHIS TagAntibody HIS.H8 (Invitrogen Co.) and StrepMAB-Classic, HRP conjugate (IBA Co.) respectively.

### Split luciferase complementation assay

*N. benthamiana* leaves were infiltrated with *A. tumefaciens* GV3101 carrying pCambia plasmids with the genes in fusion with the N-terminal or C-terminal part of luciferase (43). Three days after agroinfiltration discs were removed from the leaves using a cork borer size 2 and placed in 100 *µ*l water in a well of a 96-well plate. After a one-hour incubation a solution of 2 mM luciferin in water was added to each well and the relative light units were determined with a luminometer (Biotek Co.) over a period of 45 min. The expression level of recombinant proteins was determined by Western blotting.

### Reactive oxygen species assay

ROS assays were performed as described in (44,45). Leaf discs were removed from *N. benthamiana* leaves and incubated overnight in water, in white, flat-bottom, 96-well plates (Greiner Bio-One Co.). After 16 hours the water was replaced with a water solution containing: 34 *µ*g/ml luminol (Sigma-Aldrich Co.) and 20 *µ*g/ml horseradish peroxidase (type VI-A, Sigma-Aldrich and the designated concentration of flg22 and csp22. The relative light units were determined with a luminometer (Biotek Co.) over a period of 45 min.

### Plant growth conditions

*N. benthamiana* accession Nb-1 (46) was used in all experiments. Nb-1 plants were grown for 4-6 weeks in a controlled environment chamber with 16 hours light, 65% relative humidity and a temperature of 24°C (light period) and 22°C (dark period).

### Statistical analysis

The normality of the data was assessed using normal quantile plots (q-q plots) and the Shapiro-Wilk test. The equality of the variance was as assessed using the Bartlett test and variance of the samples is indicated by the standard deviation represented by error bars. Welch’s or Tukey-Kramer tests were used to test the differences between the means, *p* values are indicated in the Figure legends. Statistical analysis was performed with JMP Pro 14 software.

## Results

### Pti1b interacts with phosphatase PP2C6 *in vitro* and in plant cells

To identify plant proteins that interact with Pti1b, constructs encoding the kinase and an inactive variant Pti1b(K96N), both with a C-terminal 2xStrep-FLAG epitope tag, were individually introduced into leaves of *N. benthamiana* using Agrobacterium-mediated transient transformation (agroinfiltration) and Pti1b proteins were purified with FLAG-immunoprecipitation followed by Strep pull-down (Supplementary Fig. S1). Pti1b has S-acylation sites in its N terminus which play a role in its localization to the cell periphery (26). As a control, therefore, we applied the same purification method using another membrane-associated protein from Arabidopsis, PIP2A (AT3G53420) (35). The samples obtained were submitted for mass spectrometry analysis to identify potential Pti1b-interacting proteins. A protein was considered a candidate interactor if a derived peptide was detected in at least two samples with Pti1b or Pti1b(K96N) and was absent in all PIP2a samples (Fig. 1A).

**Figure 1.**
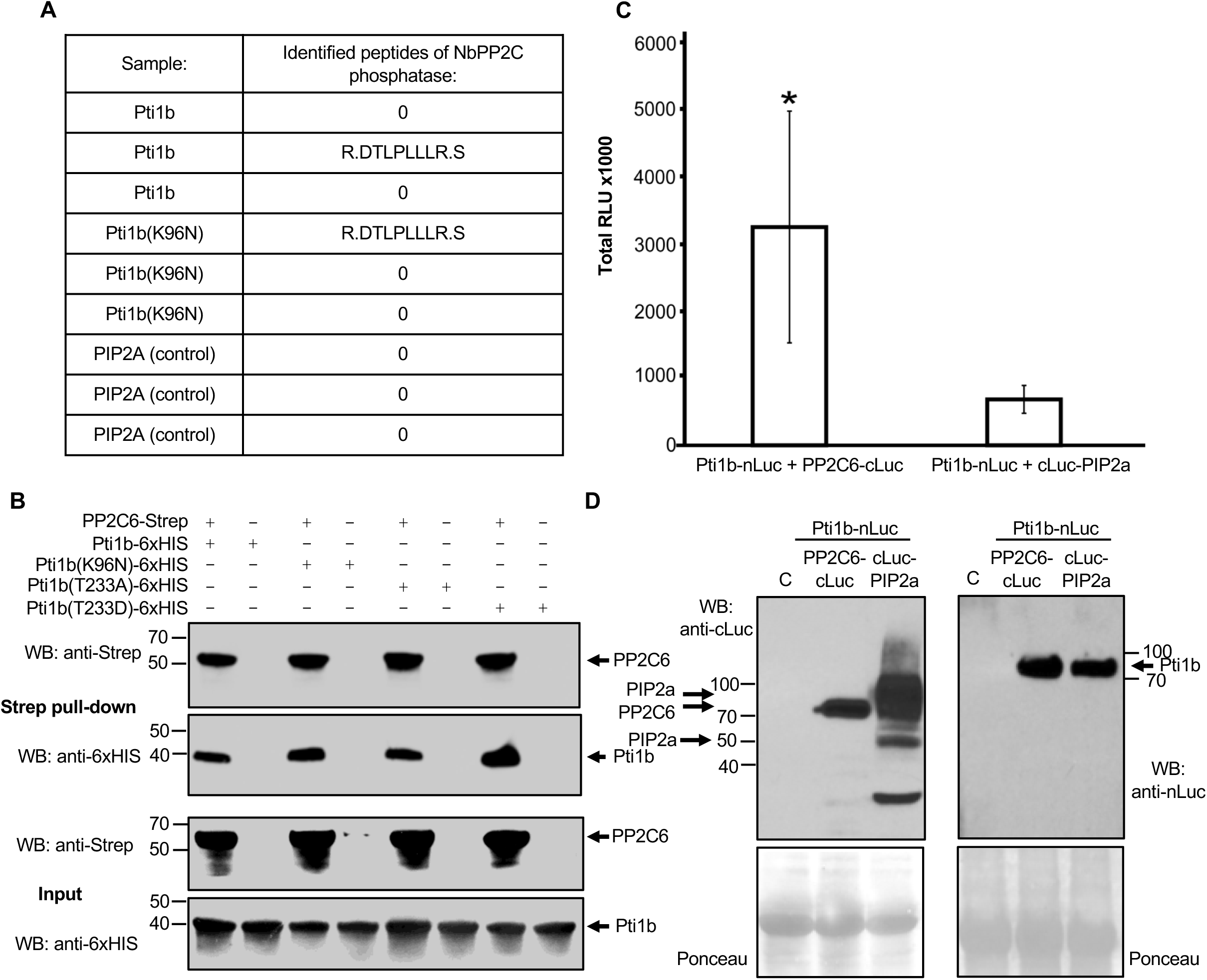
Pti1b interacts with protein phosphatase PP2C6 from tomato *in vitro* and in plant cells. (**A**) Pti1b-2xStrep-FLAG was expressed in *N. benthamiana* leaves and purified by affinity chromatography using an anti-FLAG antibody and Strep binding resin to identify Pti1b-interacting proteins. Transmembrane protein PIP2a-2xStrep-FLAG from *A. thaliana* was used as a negative control. The purified proteins were subjected to a tryptic digest and analyzed by MS/MS. Analysis of the spectra with Mascot revealed that Pti1b copurified with a protein phosphatase homologous to Arabidopsis PP2C6 phosphatase (At1g16220) and tomato PP2C6 phosphatase (Solyc07g066260) which we refer to as NbPP2C6. The table shows peptides of NbPP2C6 detected during the mass spectrometry analysis. **(B)** Variants of Pti1b interact with PP2C6 *in vitro.* Pti1b-6XHIS variants were expressed and purified from *E. coli* and then incubated with PP2C6-Strep. After incubation, PP2C6-Strep was purified using Strep affinity chromatography. Pti1b variants were detected by Western blotting (WB) using anti-6xHIS antibody and the PP2C6-Strep was detected with an anti-Strep antibody. The experiment was repeated two times with similar results. **(C)** Pti1b-nLuc interacts with PP2C6-cLuc but not with cLuc-PIP2a in a split luciferase complementation assay. Graph shows total number of relative light units (RLU) detected over 45 min after treatment of leaf discs with 1 mM luciferin. Bars represent means ± S.D. (n=6). Asterisk indicates a significant difference based on Welch’s test (p value = 0.0223). Experiment was repeated three times with similar results. (**D**) Split-luciferase fusion proteins were detected by Western blotting (WB) using an anti-LUC antibodies after transient expression in *N. benthamiana.* PIP2a monomer (lower band) and PIP2a dimer (upper band); C, sample prepared from non-infiltrated plant tissue (negative control). To confirm equal loading, the membrane was stained with Ponceau (bottom panels; the prominent protein is Rubisco).

One of the proteins that met these criteria was a protein phosphatase 2C with close similarity to tomato phosphatase Solyc07g066260. Phylogenetic analysis indicated that the closest protein in Arabidopsis to tomato phosphatase Solyc07g066260 is Arabidopsis phosphatase AtPP2C6 (At1g16220) (36), and we will refer hereafter to this tomato phosphatase as PP2C6 (Supplementary Fig. S2). Supporting a role for this phosphatase in PTI, the expression of the *PP2C6* gene is induced in tomato in response to treatment with MAMPs flgII-28 and csp22 (Supplementary Table S1).

To confirm the interaction between Pti1b and PP2C6, we first used a Strep pull-down assay in combination with wild type Pti1b, its kinase-inactive form and two variants with substitutions at threonine-233, the major autophosphorylation site in the Pti1a kinase (28). Each Pti1b protein was purified as a 6xHIS-tagged fusion, and incubated in kinase buffer with PP2C6-Strep. After incubation, PP2C6-Strep was purified with Strep-affinity chromatography and the possible presence of Pti1b-6xHIS was tested with an anti-6xHIS antibody. As shown in Fig. 1B, each of the Pti1b variants was detected only in samples containing PP2C6.

As further confirmation we examined the interaction of Pti1b and PP2C6 in plant cells using a split-luciferase complementation assay. Pti1b-nLuc and PP2C6-cLuc or Pti1b-nLuc and cLuc-AtPIP2a (as a negative control) were transiently expressed in *N. benthamiana* leaves using agroinfiltration and the leaf discs were treated with 1 mM luciferin. We detected statistically significantly more luminescence from the leaf discs expressing Ptib-nLuc and PP2C6-cLuc than from leaf discs expressing the negative control (Fig. 1C). Western blotting confirmed expression of each of the nLuc and cLuc proteins (Fig. 1D). From these experiments, we conclude that Pti1b interacts with protein phosphatase PP2C6 and, at least *in vitro*, this interaction occurs independently of Pti1b autophosphorylation.

### Pti1b autophosphorylates *in vitro* on threonine-233

In previous work we showed that the Pti1a kinase, a homolog of Pti1b, has protein kinase activity and threonine-233 in its activation loop is its major autophosphorylation site (28). The present study focused on Pti1b because a subsequent RNA-Seq experiment found that expression of the *Pti1b* gene is induced by MAMPs whereas *Pti1a* is not (Supplementary Table S1). We therefore tested Pti1b and its presumed kinase-inactive variant for kinase activity in an *in vitro* assay. Wild type Pti1b strongly autophosphorylated whereas no phosphorylation was detected with the Pti1b(K69N) variant (Fig. 2A). Next, using LC-MS/MS, we investigated whether threonine-233 is also the major phosphorylation site in Pti1b as it is in Pti1a. However, we were unable to detect the _230_LHSTR_234_ peptide generated after trypsin or pepsin digestion in either phosphorylated or unphosphorylated form.

**Figure 2.**
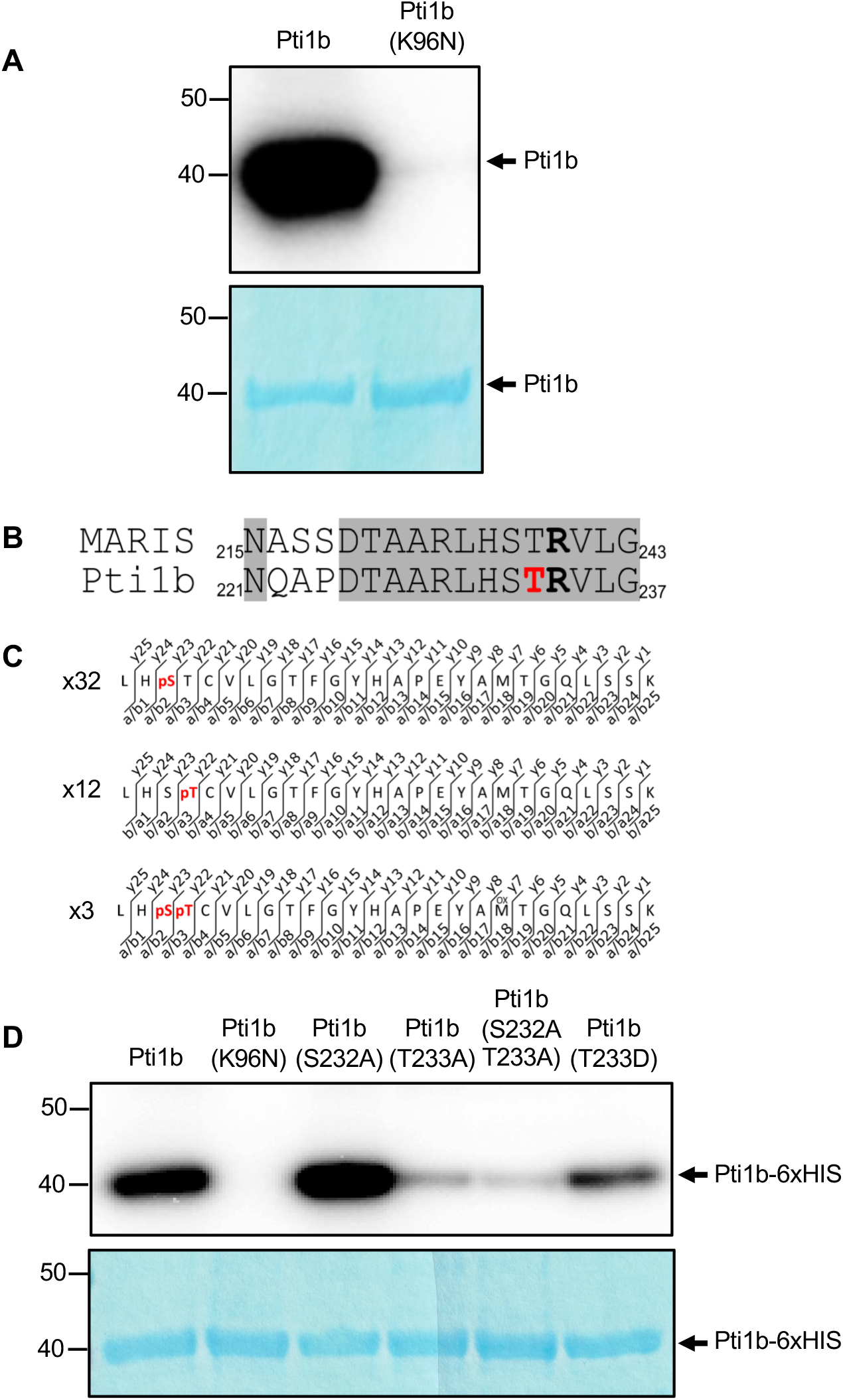
Threonine-233 is the major autophosphorylation site in Pti1b *in vitro*. **(A)** Pti1b is an active kinase with autophosphorylation capability. Pti1b and inactive variant Pti1b(K96N) were expressed and purified from *E. coli*. The proteins were incubated in kinase buffer with ATP-[γ-32] and subjected to autoradiography. The experiment was repeated three times with similar results. **(B)** Alignment of amino acid sequences of Arabidopsis MARIS and Pti1b showing the arginine-240 (bold) in MARIS and corresponding arginine-234 (bold) in Pti1b, and threonine-233, the major site of phosphorylation in Pti1b, in red; identical residues are shown in grey background. **(C)** Mass spectrometry analysis showed that variant Pti1b(R234C) is phosphorylated on serine-232 and threonine-233. Pti1b(R234C) was phosphorylated *in vitro* and the phosphorylation sites were mapped with mass spectrometry. The figure shows phosphopeptides detected during the analysis, with phosphorylated residues in red. The numbers on the left indicate the total number of times each phosphopeptide was identified in two independent experiments. Representative spectra for each phosphopeptide are shown in Supplementary Figure S3. **(D)** Threonine-233 is the major autophosphorylation site in Pti1b. Different variants of Pti1b were produced in *E. coli* and purified with affinity chromatography, and the proteins were incubated in kinase buffer with ATP-[γ-32]. The signal from phosphorylated Pti1b was detected with autoradiography and equal loading of the proteins was confirmed with Coomassie blue staining. The experiment was repeated three times with similar results.

To overcome this obstacle, we generated a Pti1b variant where the arginine-234 at the trypsin digestion site was substituted to cysteine. This variant was based on a recent report that the MARIS kinase from Arabidopsis, which belongs to the same subfamily VIII of RLCKs as Pti1b, gains constitutive function when it carries the analogous arginine-to-cysteine substitution (R240C; Fig. 2B) (18). Such a modified Pti1b, when digested with trypsin, would generate a longer peptide containing threonine-233 (LHSTCVLGTFGYHAPEYAMTGQLSSK) that might be more easily detected by LC-MS/MS. Indeed, with LC-MS/MS analysis of the autophosphorylated Pti1(R234C) we observed the presence of both phosphorylated and unphosphorylated forms of the longer peptide. Both threonine-233 and, unexpectedly, serine-232 were found to be phosphorylated in this peptide (Fig. 2C, Supplementary Fig. S3).

To investigate this observation further, we created a series of Pti1b variants with substitutions at serine-232 and threonine-233: Pti1b(S232A), Pti1b(T233A), Pti1b(S232A,T233A) and a potential phosphomimic substitution, Pti1b(T233D). These Pti1b variants were expressed in *E. coli*, purified and incubated in kinase buffer. Pti1b wild type and Pti1b(K96N) were used as a positive and negative control respectively. The results obtained with autoradiography showed that substitution of threonine-233 to either alanine or aspartic acid greatly decreased the autophosphorylation of Pti1b (Fig. 2D). In contrast, the S232A substitution had no impact on autophosphorylation of Pti1b. These data indicate that threonine-233 is likely the major site of autophosphorylation in Pti1b and suggest that the R234C substitution may cause enhanced phosphorylation of S232.

### PP2C6 dephosphorylates Pti1b *in vitro* and in plant cells

The interaction of PP2C6 with Pti1b suggests the phosphatase might act to dephosphorylate the kinase possibly as a mechanism of desensitizing its activated state. To test this possibility, we incubated Pti1b-6xHIS in kinase buffer, either alone or in the presence of PP2C6-Strep or a variant of PP2C6-Strep that has the aspartate residues at positions 110 and 234 substituted to asparagine [PP2C6(NN)-Strep]. Such asparagine substitutions are known to abolish phosphatase activity of PP2C proteins (23). We observed that in the presence of PP2C6-Strep the phosphorylated form of Pti1b was greatly reduced as compared to Pti1b incubated alone or in the presence of the inactive variant PP2C6(NN)-Strep (Fig. 3A).

**Figure 3.**
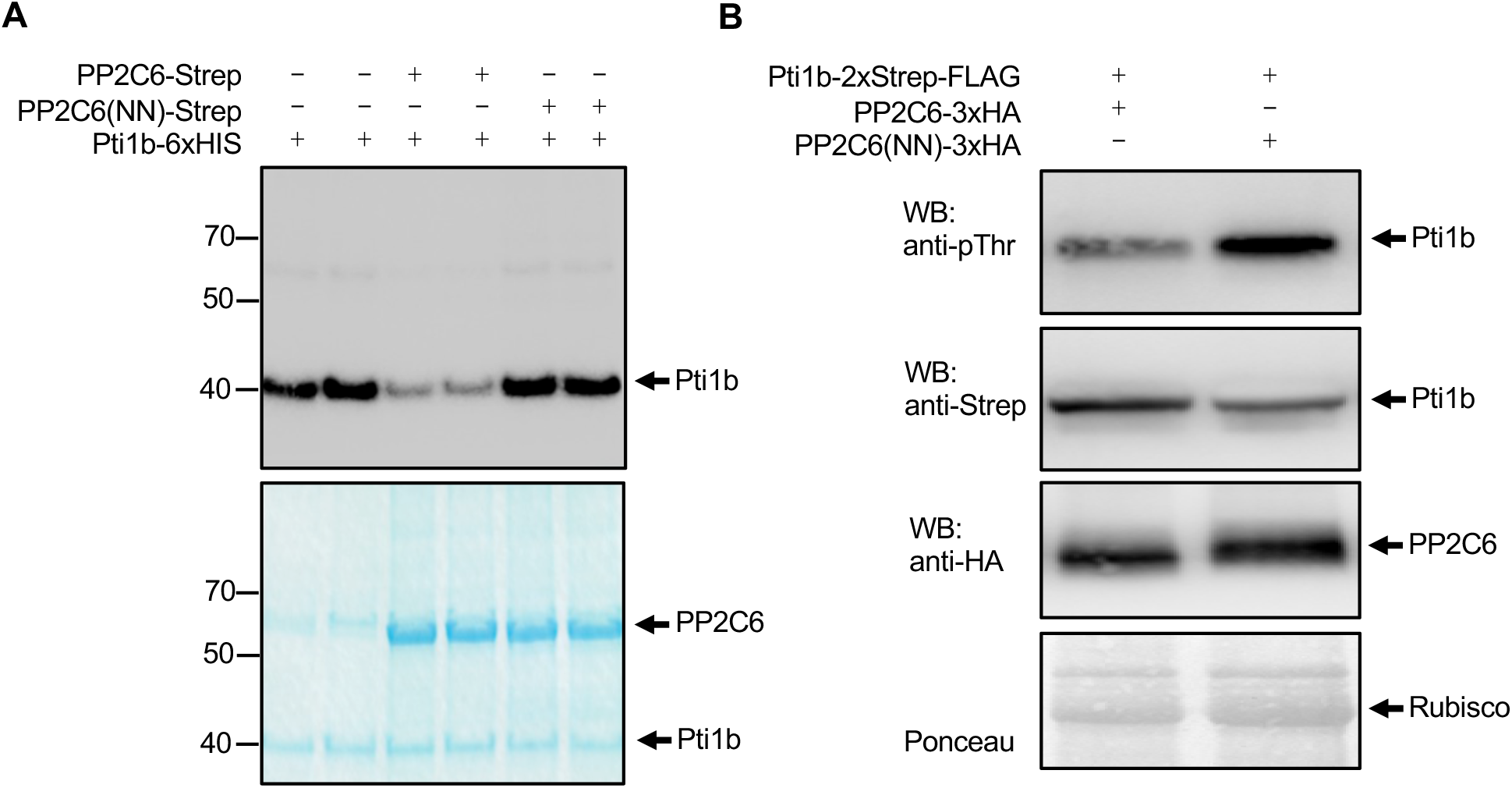
PP2C6 reduces the amount of autophosphorylated Pti1b in comparison to the catalytically-inactive variant PP2C6(NN) *in vitro* and in plant cells. (**A**) PP2C6 dephosphorylates Pti1b *in vitro*. Pti1b and PP2C6 proteins were produced in *E. coli* and purified with affinity chromatography. Pti1b was incubated alone or with the two PP2C6 variants in kinase buffer with ATP-[γ-32]. The reaction was terminated by boiling the samples in Laemmli buffer. Proteins were separated using gel electrophoresis and phosphorylated Pti1b was detected with autoradiography. Protein amounts were detected with Coomassie blue staining. Two technical repeats for each condition are shown. The experiment was repeated three times with similar results. **(B)** PP2C6 can reduce phosphorylation status of Pti1b in plant cells. Agroinfiltration was used to co-express Pti1b in leaves of *N. benthamiana* with PP2C6 or PP2C6(NN). Six days after infiltration flg22 was applied and 10 minutes later leaf tissue was harvested and Pti1b was purified using Strep-affinity chromatography. The phosphorylation level of the kinase was detected by Western blotting (WB) with anti-phosphothreonine antibodies. The total amount of Pti1b was determined with anti-Strep antibodies. The presence of PP2C6 variant in crude protein extract used for Pti1b purification was determined with anti-HA antibodies. Rubisco stained with Ponceau was used as a control of protein loading for samples with crude protein extract.

We further examined whether PP2C6 affects Pti1b phosphorylation *in vivo.* Using agroinfiltration we co-expressed Pti1b-2xStrep-FLAG with either PP2C6-3xHA or PP2C6(NN)-3xHA in leaves of *N. benthamiana*. Six days after agroinfiltration the plant tissue was treated with 1 uM flg22 for 10 minutes. Pti1b was then purified with Strep-affinity chromatography and the phosphorylation level of Pti1b was tested with antibodies that specifically recognize phosphorylated threonine residues. The specificity of the antibodies for phosphorylated Pti1b was tested with different *in vitro* phosphorylated Pti1b variants (Supplementary Fig. S4). We observed that the phosphorylation level of Pti1b-2xStrep-FLAG co-expressed with PP2C6-3xHA is reduced in comparison to Pti1b co-expressed with PP2C6(NN)-3xHA (Fig. 3B). Collectively, the data from Figs. 2 and 3 indicate that PP2C6 dephosphorylates threonine-233, the major *in vitro* autophosphorylation site in Pti1b.

### Pti1b(R234C) is recalcitrant to PP2C6 phosphatase activity *in vitro*

The arginine-to-cysteine substitution at position 240 in MARIS causes the protein to gain a constitutive-active function which restores pollen tube cell wall integrity in an *anx1*/*anx2* Arabidopsis mutant (18). The underlying reason for this gain-of-function activity is unknown (23,37). The corresponding arginine in Pti1b, at position 234, is adjacent to threonine-233 and we speculated that a R234C substitution might impact the kinase activity of Pti1b. To test this possibility, we compared the autophosphorylation activity of Pti1b and Pti1b(R234C) in an *in vitro* kinase assay and found no difference between them (Fig. 4A). We next examined whether the R234 substitution might impact the activity of PP2C6 towards Pti1b. The kinase activity of Pti1b and Pti1b(R234C) were assayed in the presence of PP2C6, PP2C6(NN), calf intestinal phosphatase (CIP) or shrimp alkaline phosphatase (rSAP). Remarkably, PP2C6 dephosphorylated Pt1b but was incapable of dephosphorylating Pti1b(R234C) (Fig. 4B). Both Pti1b proteins were effectively dephosphorylated by CIP and rSAP (Fig. 4B); for unknown reasons, CIP was phosphorylated independently of Pti1b (Supplementary Fig. S5). We next investigated whether the inability of PP2C to dephosphorylate Pti1b(R234C) was due to the loss of an interaction between these two proteins. Using the Strep pull-down assay described above we found that PP2C6-Strep interacted just as well with Pti1b as it did with Pti1b(R234C) (Fig. 4C). Thus, the Pti1b(R234C) variant has enzymatic activity comparable to Pti1b and still interacts with PP2C6 although it resists PP2C6 phosphatase activity.

**Figure 4.**
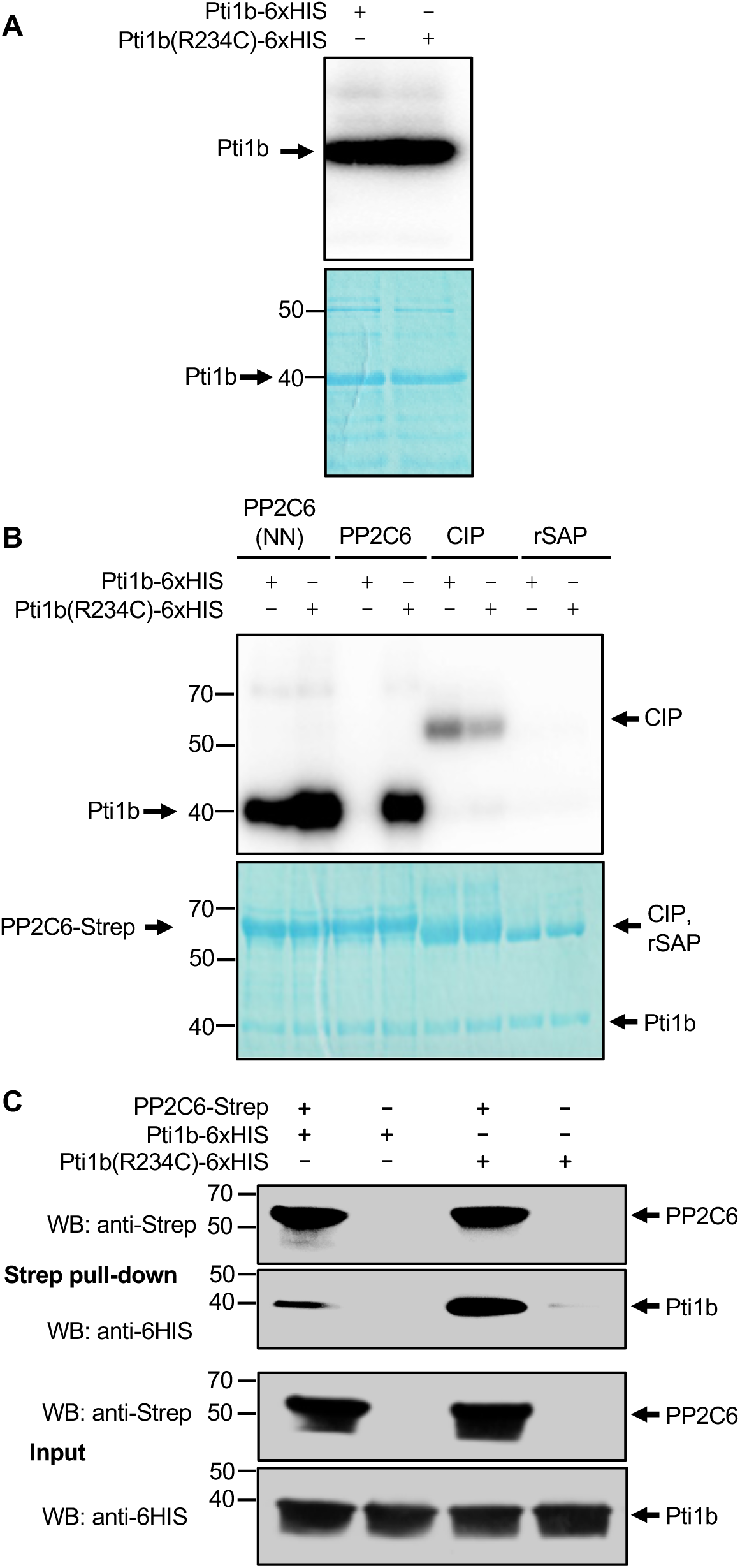
Pti1b(R234C) is recalcitrant to PP2C6 but is sensitive to other phosphatases *in vitro*. (**A**) Autophosphorylation levels of Pti1b and Pti1b(R234C) are comparable. Experiment was carried out as in the Figure 2A and experiment was repeated three times with similar results. Lower panel is Coomassie blue-stained gel. **(B**) Pti1b(R234C) is insensitive to PP2C6. Pti1b and Pti1b(R2324C) were incubated in kinase buffer in the presence of PP2C6(NN), PP2C6, CIP (calf intestinal phosphatase), or rSAP (shrimp alkaline phosphatase) and phosphorylation was detected by autoradiography. The observed phosphorylation of CIP phosphatase is not dependent on Pti1b kinase activity (Supplementary Figure S5). This experiment was repeated three times with similar results. **(C**) Pti1b(R234C) interacts with PP2C6 *in vitro.* Proteins were expressed and purified from *E. coli.* Pti1b variants were then incubated with PP2C6 phosphatase. After incubation PP2C6 phosphatase was purified with Strep-tag affinity chromatography. Pti1b variants were detected by Western blotting (WB) using an anti-6xHIS antibody and PP2C6 was detected with anti-Strep antibody. The experiment was repeated two times with similar results.

### Peptide flg22 induces the phosphorylation of Pti1b on threonine residue(s) in plant cells

Perception of MAMPs like peptide flg22 by PRRs induces phosphorylation of downstream signaling components (13,15). To test possible MAMP induction of Pti1b phosphorylation we chose to use Pti1b(R234)-Strep to avoid dephosphorylation by PP2C6. Pti1b(R234)-Strep was expressed in *N. benthamiana* leaves using agroinfiltration and the leaf tissue was infiltrated 3 days later with 1*µ*M flg22 or with water as a control. Ten minutes after addition of flg22 the tissue was collected and Pti1b(R234C) protein was purified by Strep-affinity chromatography. One half of the purified protein was then incubated with CIP to remove phosphate from Pti1b(R234C) and the samples were analyzed by Western blotting using the antibody that specifically detects phosphorylated threonine residues. This analysis revealed a higher level of phosphothreonine on Pti1b(R234C) from flg22-treated leaf tissue than from the control treated with water alone (Fig. 5). In turn, Pti1b(R234C) protein purified from the water-treated leaf tissue was phosphorylated to a higher level than the CIP-treated samples indicating that Pti1b is phosphorylated at a basal level even without flg22 treatment.

**Figure 5.**
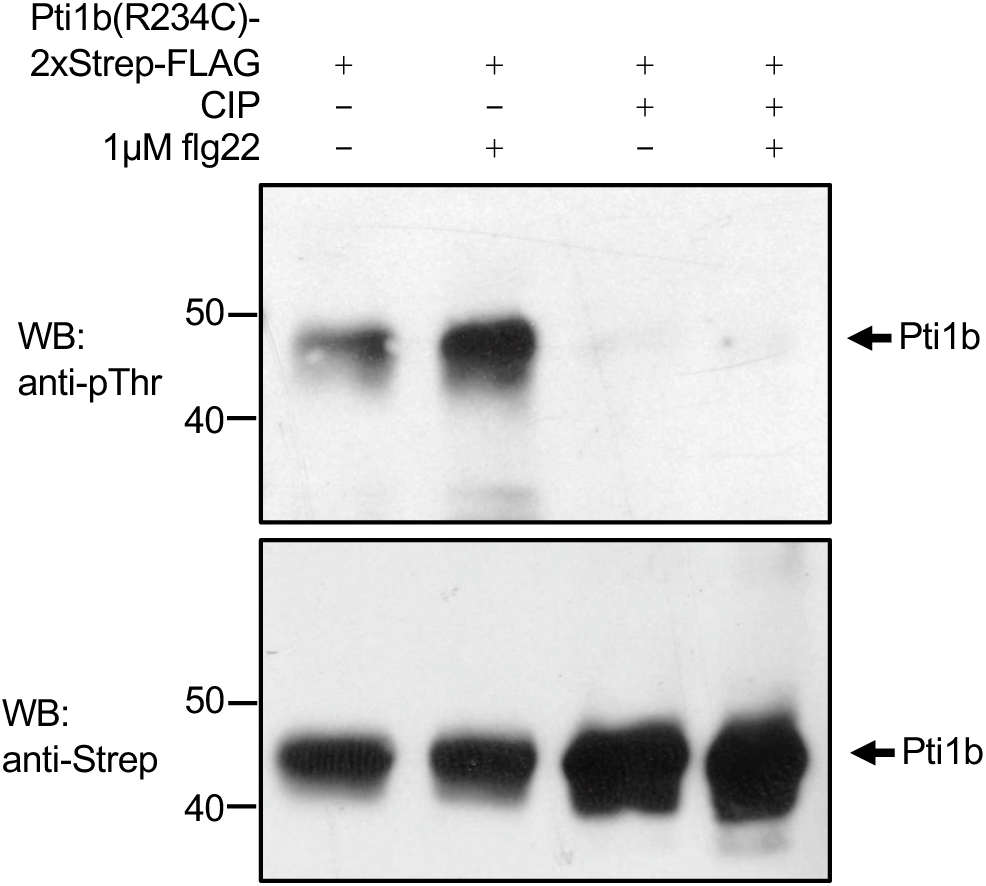
Peptide flg22 induces Pti1b(R234C) phosphorylation on threonine residue(s) in plant cells. Pti1b(R234C)-2xStrep-FLAG was expressed in *N. benthamian*a leaves using Agrobacterium-mediated transient expression (agroinfiltration). Three days after agroinfiltration, leaf tissue was infiltrated with 1µM flg22 or water as a control. Ten minutes after infiltration the tissue was collected and frozen in liquid nitrogen. Pti1b was purified with Strep-affinity chromatography. Half of the obtained samples were treated with calf intestine phosphatase (CIP) and half were untreated. Pti1b phosphorylation was detected by Western blotting (WB) with antibodies specific to phosphorylated threonine. Protein loading was tested using anti-Strep antibody. This experiment was repeated two times with similar results.

### PP2C6 decreases the production of reactive oxygen species generated in leaves in response to flg22 and csp22

Tomato plants with decreased levels of Pti1a and Pti1b produce less ROS after treatment with flg22 compared to wild type plants (26). We hypothesized that the dephosphorylation of Pti1b by PP2C6 might negatively regulate the role of Pti1 in generating ROS in response to flg22. To test this hypothesis, we used agroinfiltration to overexpress PP2C6-3xHA in leaves of *N. benthamiana*, which has four Pti1 homologs (26), and measured the production of ROS in response to flg22 and another MAMP, csp22. As controls, we overexpressed the inactive PP2C6(NN)-3xHA variant. Leaf discs expressing PP2C6 produced statistically significantly lower levels of ROS after treatment with flg22 and csp22 compared to leaf disks expressing PP2C6(NN)-3xHA (Fig. 6AC). Each of the proteins was shown to be expressed well by Western blotting (Fig. 6BD). PP2C6 therefore acts as a negative regulator of ROS production in plant leaves, likely by dephosphorylating Pti1b.

**Figure 6.**
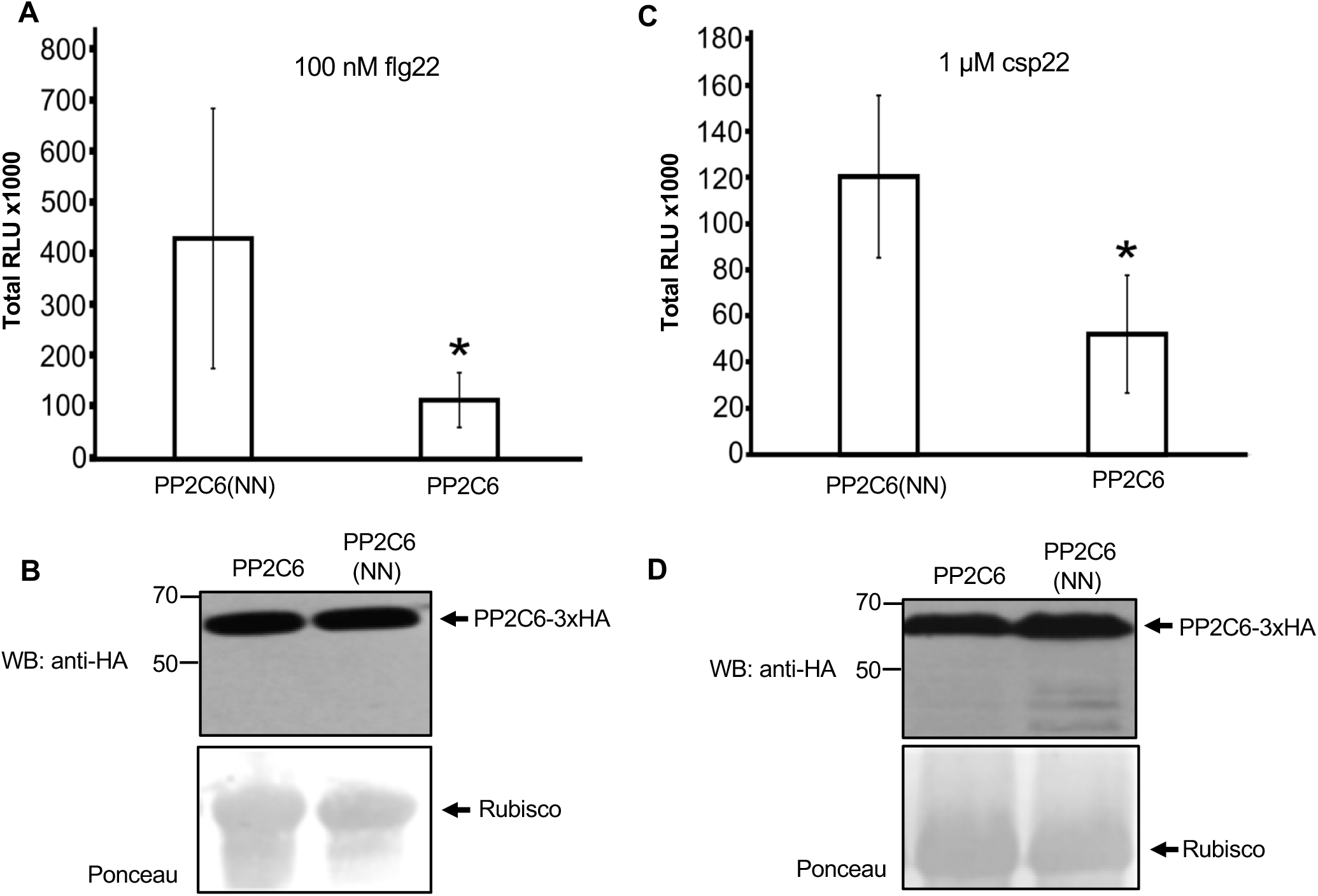
PP2C6 reduces the amount of reactive oxygen species (ROS) produced in *N. benthamiana* leaves in response to MAMPs. PP2C6 and PP2C6(NN) were expressed in *N. benthamiana* leaves using agroinfiltration, leaf disks were excised and treated with flg22 or csp22. (**A**) Total number of relative light units (RLU) detected over 45 minutes after treatment of leaf discs with 100 nM flg22. Bars represent means ± S.D. (n=8). Asterisk indicates a significant difference based on Welch’s test (p = 0.0095). **(B)** PP2C6 variants analyzed in A were detected in leaf tissue by Western blotting (WB) using an anti-HA antibody. **(C)** Total number of relative light units (RLU) detected over 45 minutes after treatment of leaf discs with 1 uM csp22. Bars represent means ± S.D. (n=8). Asterisk indicates significant difference based on Welch’s test (p = 0.0009). (**D)** PP2C6 variants analyzed in C were detected in leaf tissue by Western blotting (WB). To confirm equal loading, the membrane was stained with Ponceau (bottom panels). All the experiments were repeated two times with similar results.

## Discussion

We reported previously that the Pti1 kinases are activators of ROS burst in tomato and positive regulators of disease resistance against *P. syringae* pv. tomato (26). Here we identified PP2C6 as an interactor of Pti1 with the ability to dephosphorylate the major Pti1b autophosphorylation site, threonine-233. Phosphorylation of Pti1b is induced upon treatment of leaves with MAMPs and overexpression of PP2C6 in leaves greatly reduces production of immunity-associated ROS. Below we elaborate on our key findings about Pti1b phosphorylation status, the role of PP2C6 in regulating Pti1b, and on the ability of the arginine-234-cysteine substitution to make Pti1b recalcitrant to PP2C6 activity.

### Pti1b phosphorylation

Our mass spectrometry experiments suggested that Pti1b is autophosphorylated *in vitro* on threonine-233 and/or on serine-232. However, with autoradiography we were only able to find evidence for Pti1b phosphorylation on threonine-233 and our current data therefore supports this residue as the major phosphorylation site. Phosphorylation of serine-232 was also not observed in a previous study that reported threonine-233 as the major site of phosphorylation in the Pti1a kinase (28). Interestingly, we do observe weak phosphorylation associated with both Pti1b(T233A) and Pti1b(T233D). The source of this signal could be phosphorylated serine-232 which may be a minor site of autophosphorylation in Pti1b. It is possible that the arginine-234-cysteine substitution causes more phosphorylation to occur on serine-232 than occurs in wildtype Pti1b. This possibility is supported by the greater number of peptides observed by mass spectrometry with phosphorylated S232 (Fig. 2C) and by the experiment in which we compared the autophosphorylation of Pti1b(T233A) and Pti1b(T233A/R234C) and found the latter protein to be more strongly phosphorylated (Supplementary Fig. S4A). It also appears that the serine-232-alanine substitution might strengthen the autophosphorylation on threonine-233 as shown in Fig. 2B.

We found that flg22 induces the phosphorylation of Pti1b on threonine residue(s) in *N. benthamiana.* Although we have not investigated this further yet, it is likely this phosphorylation occurs on threonine-233 because that residue is the major phosphorylation site *in vitro*. It has been reported previously that OsPti1a is phosphorylated in rice cells on its threonine-233 (corresponding to threonine-233 in Pti1b) (38). Additionally, it was shown that threonine-233 of OsPti1a plays a role in immunity against *X. o.* pv. oryzae (39). Further insights into which residues of Pti1b are phosphorylated in plant cells will come from the future development and biochemical characterization of transgenic tomato plants with phosphonull and phosphomimic variants of threonine-233.

### The role of phosphatase PP2C6

There are relatively few protein phosphatases which are known regulate RLCKs that are activators of plant immune responses. One of these, PP2C38 (23), which regulates the immunity-associated RLCK BIK1, has some similarities to PP2C6. Like PP2C6 and Pti1b, PP2C38 negatively controls the phosphorylation status of BIK1 (23). Treatment of Arabidopsis leaves with flg22 induces BIK1 activity causing it to phosphorylate PP2C38 which leads to dissociation of the phosphatase from BIK1 (23). The dissociated BIK1 is then able to activate downstream targets. We did not observe Pti1b phosphorylation of either PP2C6 or PP2C6(NN) *in vitro*. Therefore, we have no evidence that release of Pti1b from interaction with PP2C6 relieves negative regulation by PP2C6 indicating Pti1b-PP2C6 may use a different mechanism than BIK1-PP2C38. A hypothesis that would be interesting to explore is that Pti1b might be phosphorylated by an upstream kinase such as Oxi1 which allows Pti1b to mitigate the negative regulation imposed by PP2C6.

### The influence of the arginine-234-cysteine substitution on PP2C6 activity

The MARIS protein, which is a member of the same RLCK subfamily as Pti1b, acts as a regulator of cell wall integrity in Arabidopsis (18). The variant of this kinase with an arginine-to-cysteine substitution was identified as a constitutive-active version of MARIS but the underlying reason for this activity is unknown (18). Here we found that the analogous substitution in Pti1b (R234C) makes the kinase insensitive to dephosphorylation by PP2C6 although its kinase activity is comparable to Pti1b wild type. Based on this observation, it is possible that MARIS(R240C) is also able to resist dephosphorylation by an associated phosphatase that normally negatively regulates MARIS. This could explain how the constitutively-active variant of MARIS is able to restore pollen tube cell wall integrity in the absence of Anx1/Anx2.

### A model for Pti1b and PP2C6

Based on the results presented here we propose a model for Pti1b and PP2C6 in pattern-triggered immunity (Supplementary Fig. S6). Various MAMPs including flg22 and csp22 activate PRRs by binding to their extracellular leucine-rich repeat domains. These PRRs act in concert with the co-receptor BAK1 (SERK3B) in signaling pathway(s) which lead to multiple defense responses including production of ROS. Pti1b is proposed to act downstream or with PRR signaling complexes and plays a role in ROS production. Phosphorylation of threonine-233 in Pti1b is proposed to contribute to this mechanism and PP2C6 acts to dephosphorylate this residue thereby inactivating Pti1b and desensitizing the associated signaling pathway. In the future we will investigate the possibility that Pti1b is present in PRR complexes, how it might be impacted by activated PRRs, and search for its potential substrates which are expected to include proteins involved in ROS production.

## Abbreviations

ANXR: anxur receptor-like kinase
BAK1: brassinosteroid insensitive 1-associated kinase 1
BIK1: botrytis-induced kinase1
CERK1: chitin elicitor receptor kinase 1
CORE: Cold shock protein receptor
csp: cold shock protein
CWI: cell wall integrity
EGF: epidermal growth factor
flg: flagellin
FLS2: flagellin sensing 2
KAPP: kinase associated protein phosphatase
LRR: leucine-rich repeat
MAMPs: microbe-associated molecular patterns
MAPKs: mitogen-activated protein kinases
NADPH: nicotinamide adenine dinucleotide phosphate
Oxi1: oxidative signal-inducible1
PBL: avrPphB susceptible-like
PP2C: protein phosphatase 2C
PRRs: pattern recognition receptors
PTI: pattern-triggered immunity
Pti1b: Pto interactor 1b
RBOHD: respiratory burst oxidase homolog protein D
RLCKs: receptor-like cytoplasmic kinases
ROS: reactive oxygen species
XB15: XA21 binding protein 15.

## Acknowledgments

We thank Fabio Rinaldi and Robyn Roberts helpful comments on the manuscript, Konrad Thorner, Sam Wolfe, and Lydia Zamidar for technical assistance, and Gitta Coaker for the cLuc-PIP2A plasmid. This research was supported by National Science Foundation grant nos. IOS-1451754 and IOS-1546625 (GBM).

## Author contributions

FG and GBM conceived and designed the experiments, FG performed experiments, FG and GBM analyzed the data and wrote the paper

## Supporting Information

**Figure S1.**
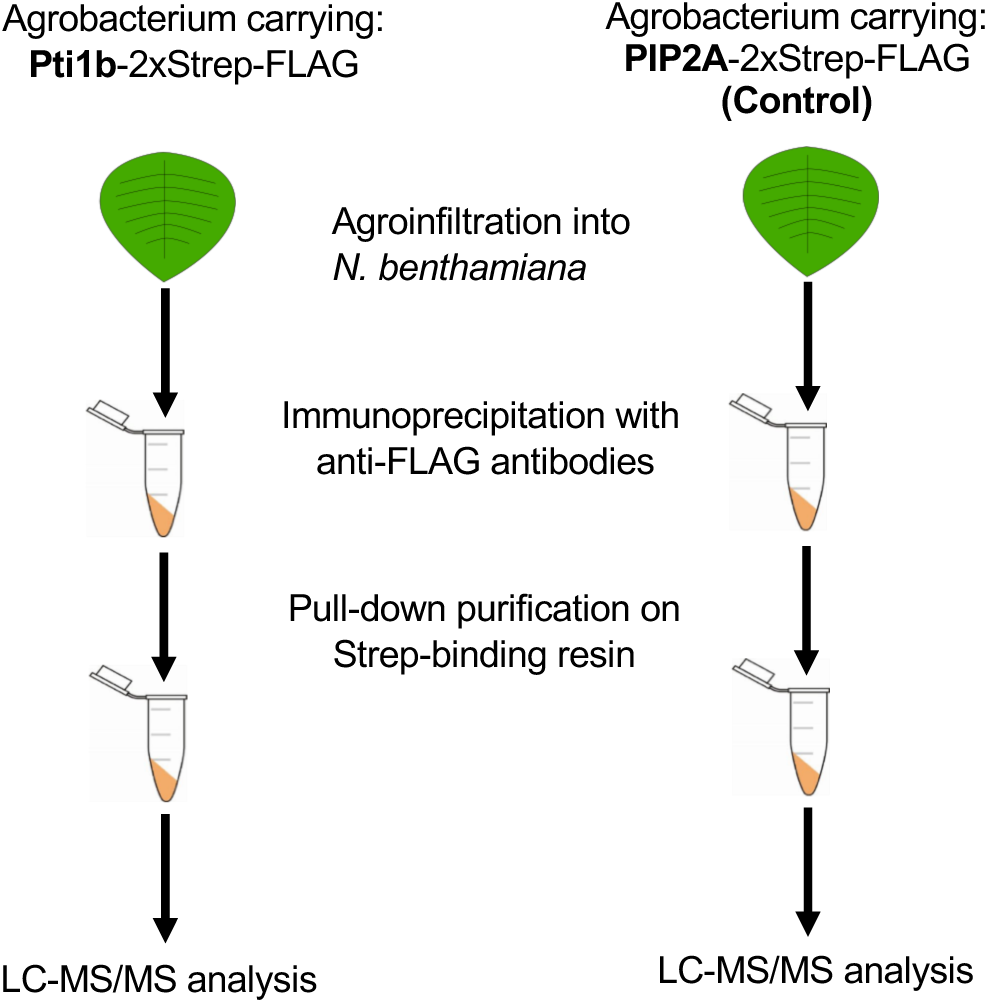
Scheme used to identify plant proteins that co-purify with Pti1b. Pti1b was expressed by agroinfiltration in *Nicotiana benthamiana* leaves and purified as shown using combined FLAG-tag and Strep-tag affinity chromatography. Transmembrane protein PIP2a was used as a control. After purification, samples were submitted for mass spectrometry analysis to detect proteins that specifically co-purified with Pti1b.

**Figure S2.**
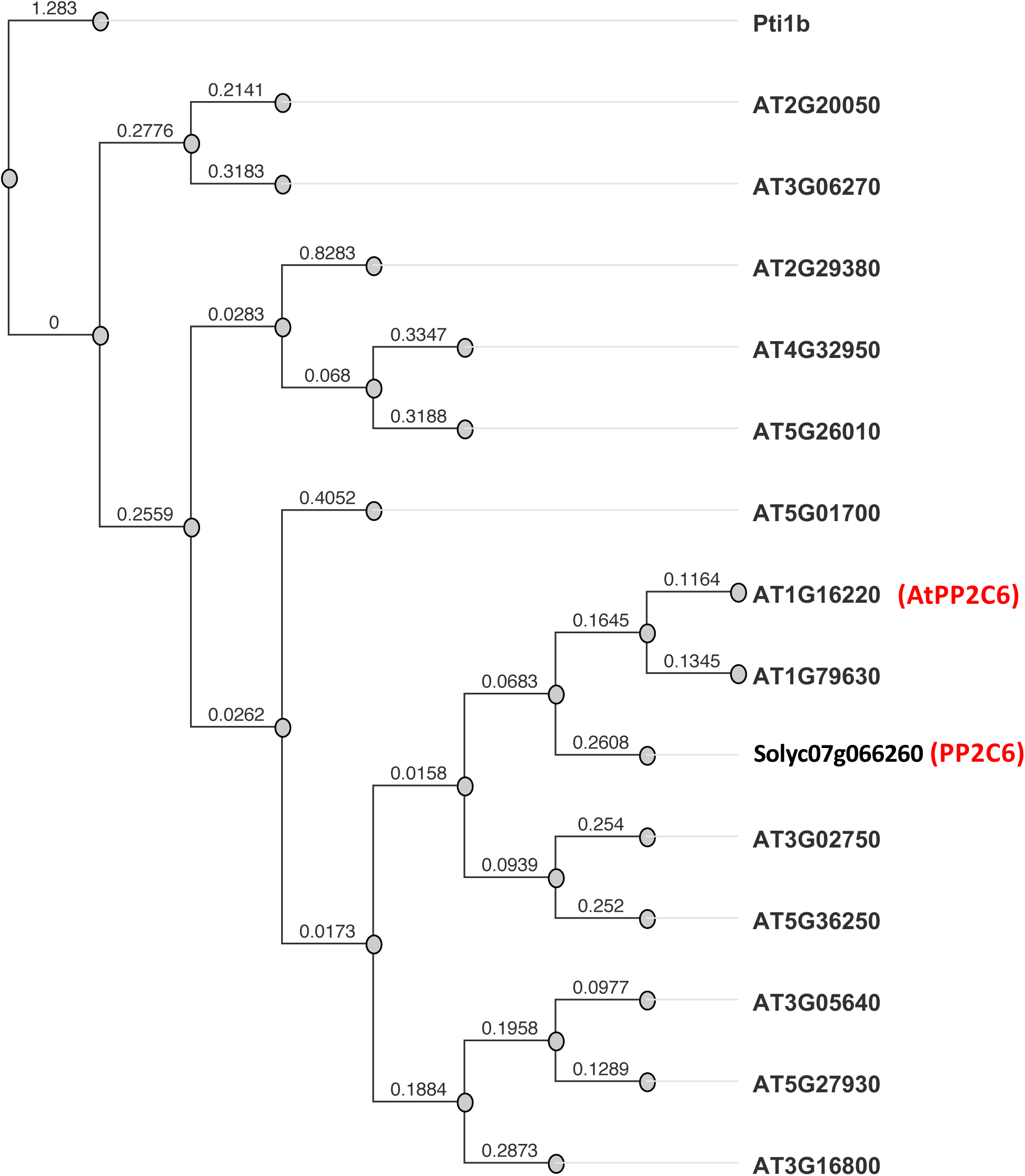
Phylogenetic tree of Arabidopsis phosphatase proteins showing relatedness of AtPP2C6 and PP2C6. The phylogenetic tree was generated with Geneious Tree Builder with the following parameters: alignment type - global alignment, cost matrix - Pam160, genetic distance model - jukes cantor, tree-build method - neighbor-joining, with the amino acid sequence of Pti1b used as the outgroup. Numbers refer to the evolutionary distances between proteins. AtPP2C6 (At1g16220) is the closest homolog of PP2C6 (Solyc07g066260).

**Figure S3.**
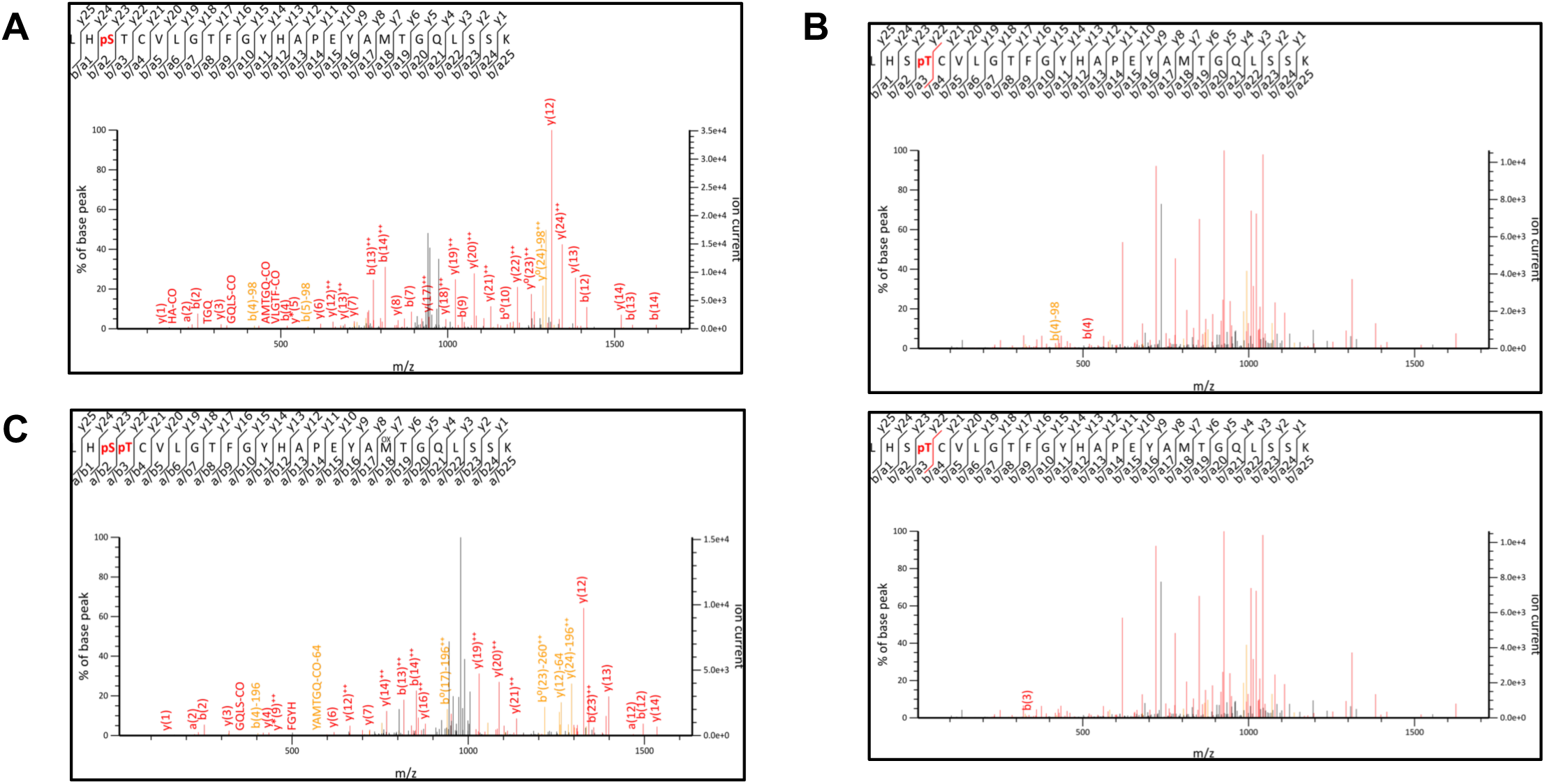
MS/MS spectra of phosphopeptides in Pti1b phosphorylated *in vitro*. **A)** Representative spectra for the peptide with phosphorylated serine-232. The ions b(4), b(5) and y(24) shown in orange are the product ions generated by loss of 98 Da (H_3_PO4) from parent ions b(4), b(5) and y(24) shown in red. **B**) Upper panel: Representative spectra for peptide with phosphorylated threonine-233. The ion b(4) shown in orange is a product of ion generated by loss of 98 Da (H_3_PO4) from parent ion b(4) shown in red. Bottom panel – the product ion of parent ion b(3) was not detected which supports the conclusion that the phosphothreonine-233 but not phosphoserine-S232 is a source of H_3_PO4 group in ion b(4) from upper panel. **C)** The ions b(4), b(5) and y(24) in orange are the product ions generated by loss of 196 Da (two H_3_PO4 group) from parent ions b(4), b(5) and y(24) in red.

**Figure S4.**
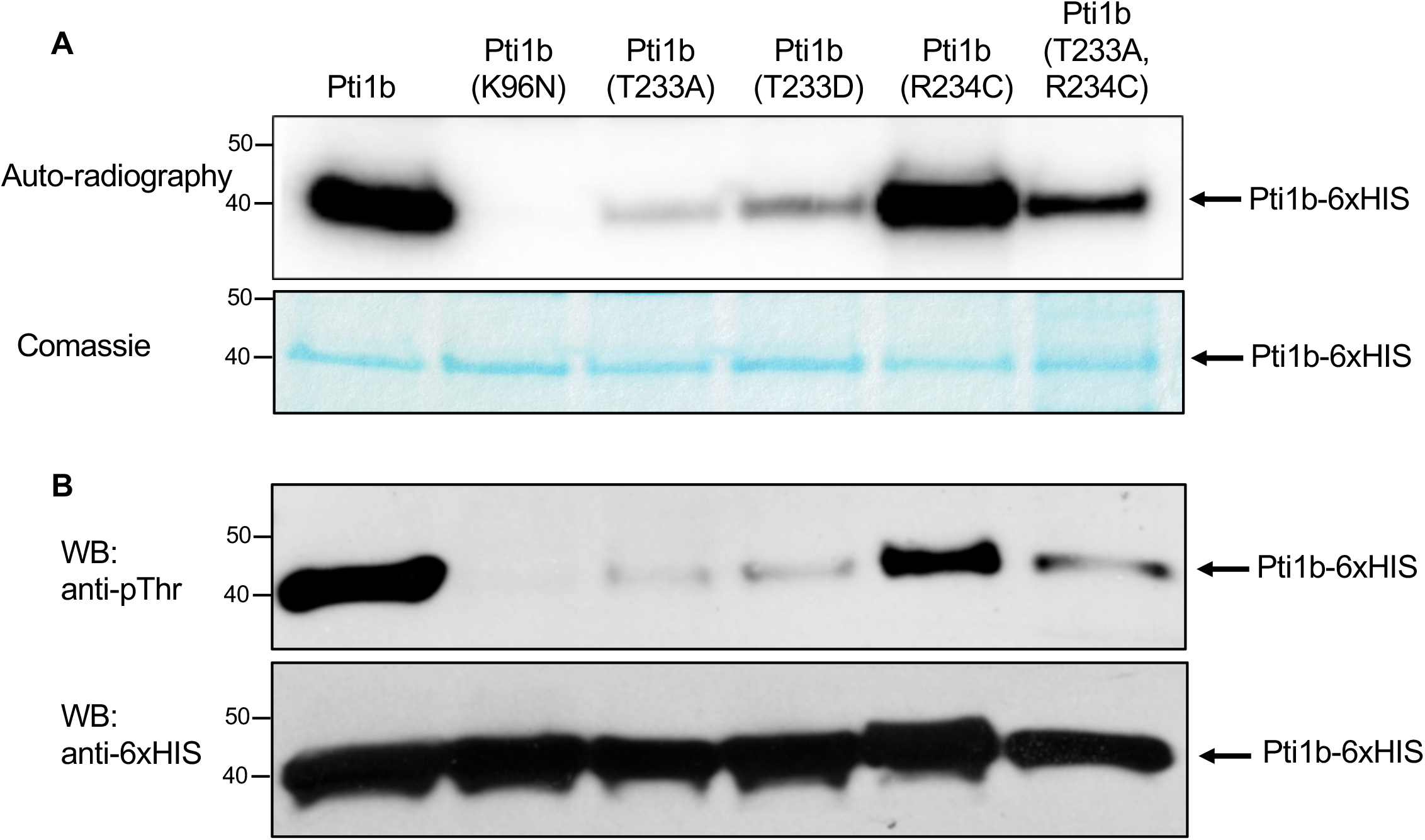
Testing the specificity of the anti-phosphothreonine antibodies used in experiment showed in Figure 5. **A)** Upper panel: autophosphorylation of different Pti1b isoforms was examined with autoradiography; lower panel: total amount of protein loaded was checked with Coomassie blue staining. **B)** Upper panel: autophosphorylation of different Pti1b isoforms was detected using Western blotting with anti-phosphothreonine antibodies; lower panel: the amount of loaded protein was tested with anti-6xHIS antibodies. The experiment was repeated twice with similar results.

**Figure S5.**
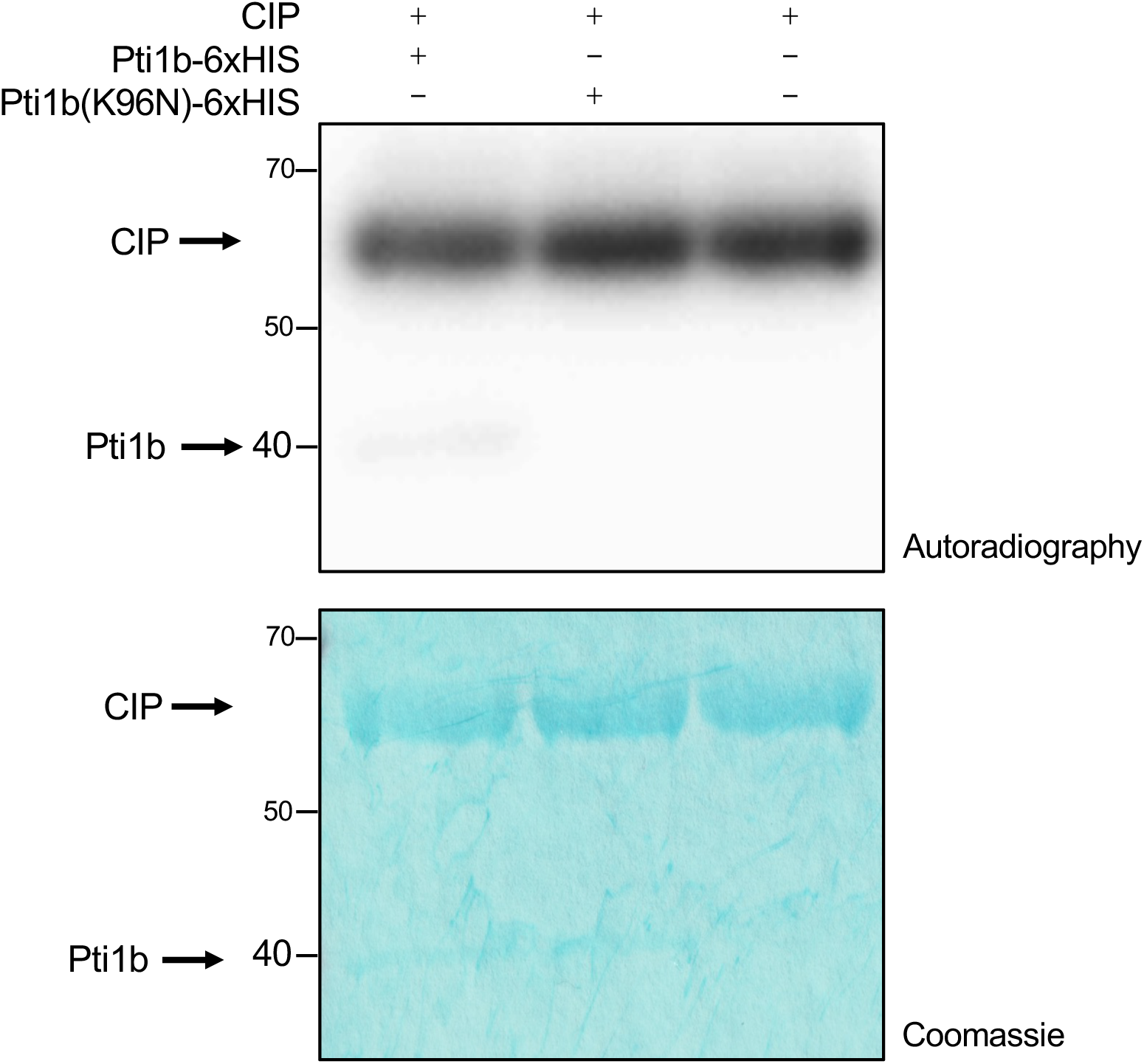
The phosphorylation status of CIP is not dependent on kinase activity of Pti1b. CIP phosphatase was incubated alone or in the presence of Pti1b or an inactive variant, Pti1b(K96N). The phosphorylation status of the proteins was examined with autoradiography (upper panel). The amount of protein loaded was checked with Coomassie staining (lower panel).

**Figure S6.**
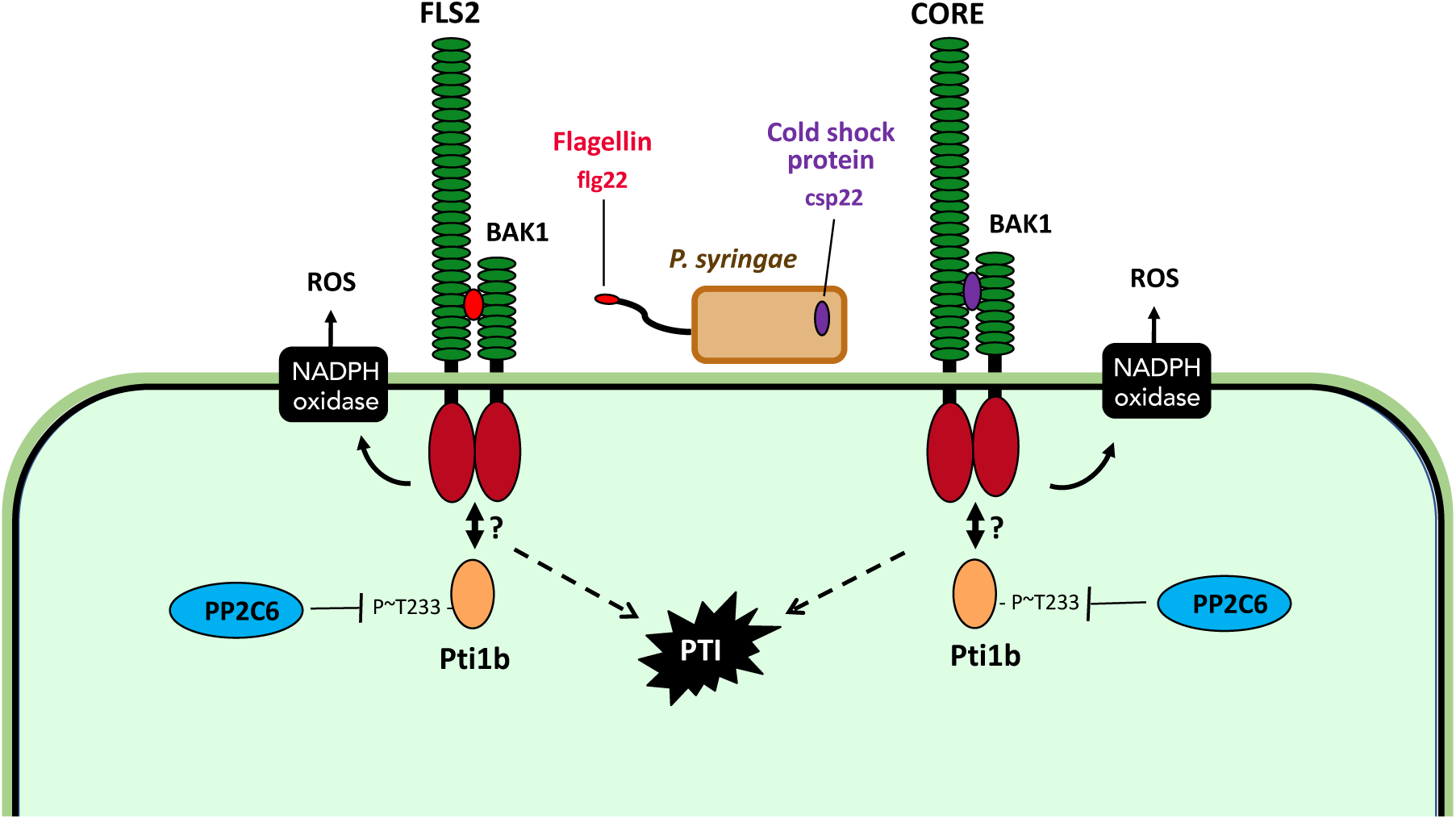
A model for negative regulation of Pti1b kinase by the PP2C6 phosphatase. The model shows that in the presence of a microbe expressing flg22 or csp22, the FLS2 and CORE receptors, respectively, directly or indirectly activate the Pti1b kinase by increasing phosphorylation of threonine-233 in Pti1b which activates downstream pattern-triggered immunity (PTI) responses. PP2C6 dephosphorylates threonine-233 in Pti1b thereby negatively regulating Pti1b-mediated PTI responses. See text for further discussion of the model.

**Table S1.** Expression of tomato *Pti1a,Pti1b*,and *PP2C6* genes in response to MAMPs. Expression of the indicated genes in response to flgII-28 or csp22 treatment of tomato leaves. RNA sequencing data are from Rosli *et al*. (2013) *Genome Biology* 14:R139 and Pombo *et al*. (2014) *Genome Biology* 15:492 and are available from TGFD (http://ted.bti.cornell.edu/cgi-bin/TFGD/digital/home.cgi, D007 and D010). Transcript abundance was measured as RPKM (reads per kilobase per exon model per million mapped reads) 6 hr after syringe-infiltration with 1 μM flgII-28, 1 μM csp22 or a buffer-only (mock) solution into Rio Grande-prf3 (carrying a mutation in *Prf*). Results shown are the ratios between the means of different treatments (*n* = 3 biological replicates), and significant differences were determined by FDR-adjusted *p*-values. Red shading indicates genes with increased transcript abundance as a result of the treatment, and green indicates no statistically significant change in transcript abundance in response to MAMP treatment.

| Gene name | Solyc no. | PTI Induction |  | PTI Induction |  |
| --- | --- | --- | --- | --- | --- |
|  |  | flgII-28/<br>mock |  | csp22/<br>mock |  |
|  |  | Ratio | FDR | Ratio | FDR |
| *Pti1a* | Solyc12g098980 | 1.07 | 1 | 1.34 | 0.4878 |
| *Pti1b* | Solyc05g053230 | 2.68 | 2.42e-07 | 2.56 | 2.31e-05 |
| *PP2C6* | Solyc07g066260 | 2.68 | 5.02e-07 | 3.28 | 8.94e-08 |

**Table S2.**
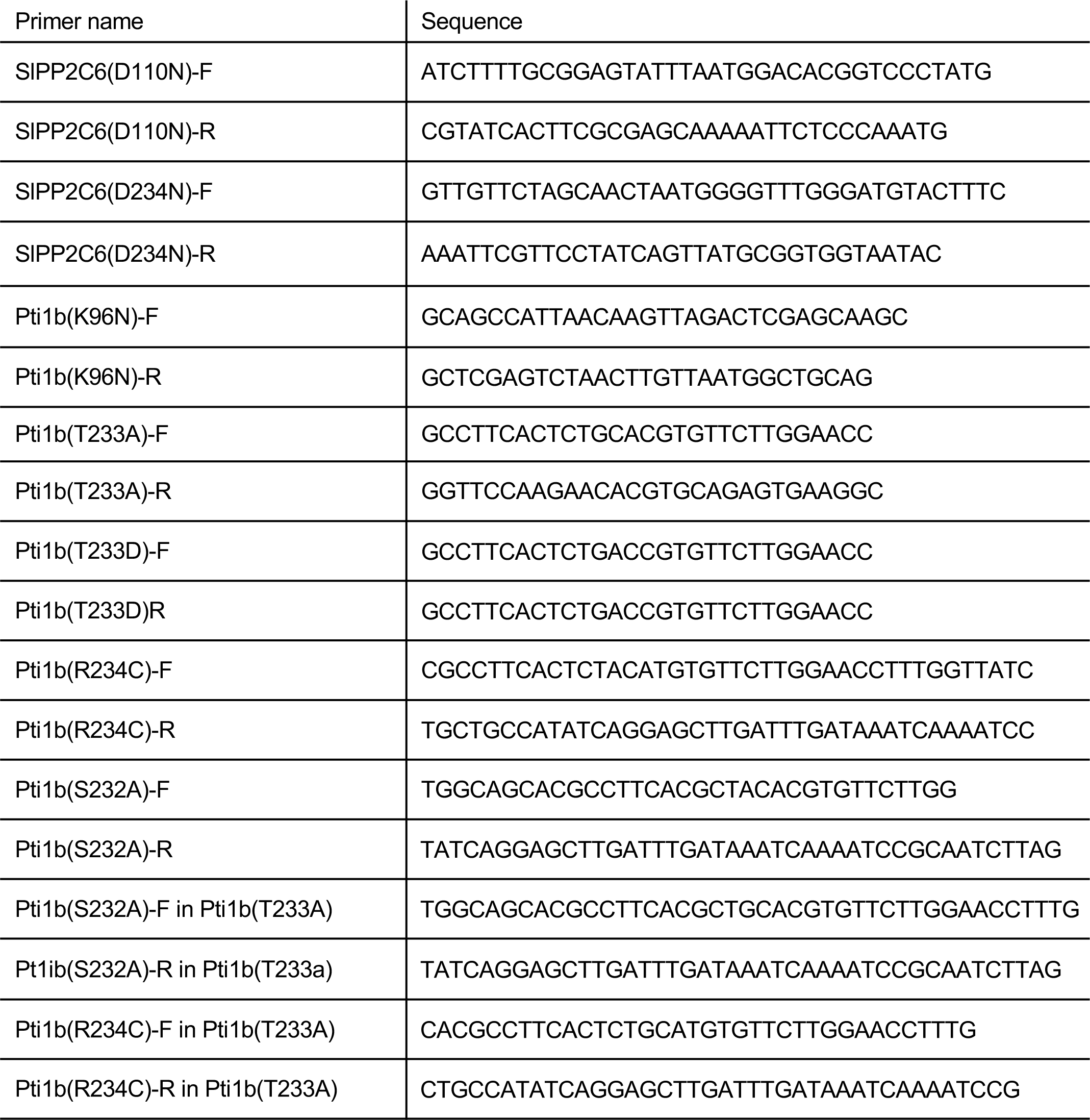
Primers used in site-directed mutagenesis.

**Table S3.**
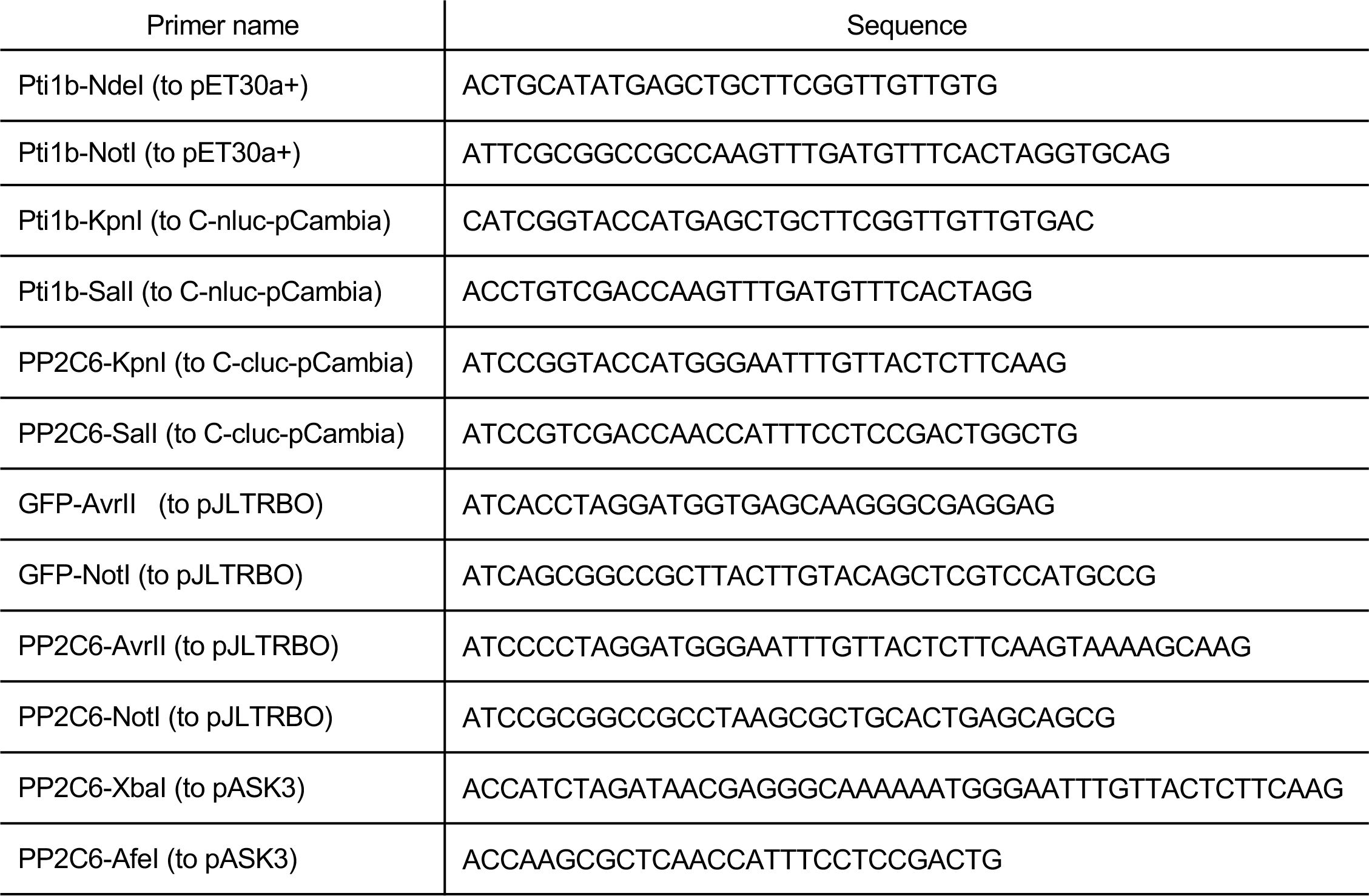
Primers used in molecular cloning.

